# Cancer causes metabolic perturbations associated with reduced insulin-stimulated glucose uptake in peripheral tissues and impaired muscle microvascular perfusion

**DOI:** 10.1101/734764

**Authors:** Xiuqing Han, Steffen H. Raun, Michala Carlsson, Kim A. Sjøberg, Carlos Henriquez-Olguín, Mona Ali, Annemarie Lundsgaard, Andreas M. Fritzen, Lisbeth L. V. Møller, Zhen Li, Jinwen Li, Thomas E. Jensen, Bente Kiens, Lykke Sylow

## Abstract

**Background:** Redirecting glucose from skeletal muscle and adipose tissue, likely benefits the tumor’s energy demand to support tumor growth, as cancer patients with type 2 diabetes have 30% increased mortality rates. The aim of this study was to elucidate tissue-specific contributions and molecular mechanisms underlying cancer-induced metabolic perturbations.

**Methods:** Glucose uptake in skeletal muscle and white adipose tissue (WAT), as well as hepatic glucose production, were determined in control and Lewis lung carcinoma (LLC) tumor-bearing C57BL/6 mice using isotopic tracers. Skeletal muscle microvascular perfusion was analyzed via a real-time contrast-enhanced ultrasound technique. Finally, the role of fatty acid turnover on glycemic control was determined by treating tumor-bearing insulin-resistant mice with nicotinic acid or etomoxir.

**Results:** LLC tumor-bearing mice displayed reduced insulin-induced blood-glucose-lowering and glucose intolerance, which was restored by etomoxir or nicotinic acid. Insulin-stimulated glucose uptake was 30-40% reduced in skeletal muscle and WAT of mice carrying large tumors. Despite compromised glucose uptake, tumor-bearing mice displayed upregulated insulin-stimulated phosphorylation of TBC1D4^Thr642^ (+18%), AKT^Ser474^ (+65%), and AKT^Thr309^ (+86%) in muscle. Insulin caused a 70% increase in muscle microvascular perfusion in control mice, which was abolished in tumor-bearing mice. Additionally, tumor-bearing mice displayed increased (+45%) basal (not insulin-stimulated) hepatic glucose production.

**Conclusions:** Cancer can result in marked perturbations on at least six metabolically essential functions; i) insulin’s blood-glucose-lowering effect, ii) glucose tolerance, iii) skeletal muscle and WAT insulin-stimulated glucose uptake, iv) intramyocellular insulin signaling, v) muscle microvascular perfusion, and vi) basal hepatic glucose production in mice. The mechanism causing cancer-induced insulin resistance may relate to fatty acid metabolism.

## 1.1. Introduction

Epidemiological and clinical studies show an association between several types of cancers and poor glycemic control in humans. For example, one-third of cancer patients are glucose intolerant [1], which is of clinical relevance as cancer patients with type 2 diabetes have 30% increased mortality rates [2, 3]. Furthermore, recent studies have suggested that insulin resistance could be an underlying cause of cancer-associated loss of muscle and fat mass, called cancer cachexia [4–6]. Cachexia occurs in 60-80% of cancer patients and is associated with poor prognosis [7]. Moreover, glucose uptake was significantly lower in lung cancer patients with cachexia during hyperinsulinemia [8]. Despite the emerging link between cancer, cancer cachexia and insulin resistance, the tissue-specific mechanistic and molecular cause(s) of insulin resistance in cancer is largely unexplored.

Skeletal muscle and adipose tissue are essential for maintaining whole-body glucose homeostasis by taking up the majority of glucose in response to insulin in healthy humans [9]. Insulin promotes glucose disposal in skeletal muscle and adipose tissue by increasing microvascular perfusion as well as mediating glucose transport across the plasma membrane [10–12]. Whether insulin resistance in cancer is associated with molecular malfunctions in skeletal muscle and adipose tissue, and whether impaired muscle microvascular perfusion is a potential cause of reduced insulin-stimulated glucose uptake have, to our knowledge, not previously been determined.

Fatty acid oxidation and/or adipose tissue lipolysis are increased in many cancers [13–15]. Excessive fatty acids in the circulation can cause insulin resistance and have been suggested as a cause of insulin resistance in obesity and type 2 diabetes [16–18]. Moreover, literature suggests that excessive fatty acid turnover is a leading cause of cancer cachexia because blockade of fatty acid oxidation or suppression of lipolysis in adipose tissue prevents cachexia in tumor-bearing mice [19–21]. However, a role for altered fatty acid turnover in cancer-associated impaired glycemic regulation is unexplored.

The aim of the present investigation was to elucidate tissue-specific contributions and molecular mechanisms underlying impaired glycemic regulation in cancer.

In a mouse model of lung cancer, we found significant whole-body insulin resistance and glucose intolerance that was restored by blockage of whole-body fatty acid oxidation or adipose tissue lipolysis. Insulin resistance in tumor-bearing mice was associated with i) impaired glucose uptake in adipose tissue and skeletal muscle despite augmented muscle insulin signaling, ii) abrogated muscle microvascular perfusion in response to insulin, and iii) increased basal hepatic glucose production.

## 2. Material and methods

### 2.1. Cell culture

Lewis lung carcinoma cells (LLC, ATCC® CRL1642™) were cultured in DMEM, high glucose (Gibco #41966-029,USA) supplemented with 10% fetal bovine serum (FBS, Sigma-Aldrich #F0804, USA), 1% penicillin-streptomycin (ThermoFisher Scientific #15140122, USA) (5% CO2, 37°C). Prior to inoculation into mice, LLC cells were trypsinized and washed twice with PBS. LLC cells were suspended in PBS with a final concentration of 2.5 * 10^6^ cells/ml.

### 2.2. Animals

Female C57BL/6 (Taconic, Lille Skensved, DK) mice were group-housed at ambient temperature (21-23°C) with nesting materials and kept on a 12 h:12 h light-dark cycle with access to a standard rodent chow diet (Altromin no. 1324, Brogaarden, DK) and water *ad libitum*. At the age of 12-14 weeks, mice were randomly assigned into control (Control) or LLC tumor-bearing (LLC) groups and subcutaneously injected with 100 µl PBS with or without 2.5 * 10^5^ LLC cells into the right flank. Control and LLC tumor-bearing mice were sacrificed at two time points: 15 days (LLC Day 15) and 21-27 days (LLC Day 21-27) after tumor transplantation. Tumor volume (V) was monitored by caliper measurement and defined by V [mm^3^] = (length [mm]) × (width [mm])^2^ × 0.52 every 2 to 5 days [22]. Food intake was measured continuously every second day during the intervention in non-tumor-bearing, tumor-bearing and etomoxir-treated (described below) tumor-bearing mice. Body weight was monitored before and throughout the interventions. All experiments were approved by the Danish Animal Experimental Inspectorate (Licence: 2016-15-0201-01043).

### 2.3. Etomoxir and nicotinic acid administration

Mice housed at ambient temperature are mildly cold stressed and preferentially metabolize more lipids than carbohydrate [23], thus inhibiting fat metabolism might cause undue metabolic stress at ambient temperature and therefore we performed this part of the experiment at thermoneutrality (30°C). After 3 weeks of acclimatization and mimicking intraperitoneal (i.p) injection every other day using empty syringes, LLC transplantation was performed as described above. Half of the tumor-bearing mice and half of the non-tumor-bearing mice were i.p. injected daily with 5 mg/kg body weight ethyl-2-[6-(4-chlorophenoxy)hexyl]-oxirane-2-carboxylate (etomoxir) (Sigma-Aldrich, US), dissolved in 5% (2-Hydroxypropyl)-β-cyclodextrin solution from day 8 following tumor transplantation (LLC-Eto). The other half of the tumor-bearing mice (LLC) and non-tumor-bearing control mice (Control) were i.p. injected with 5% (2-Hydroxypropyl)-β-cyclodextrin solution. Mice were sacrificed 18 days following tumor transplantation.

The nicotinic acid-administrated mice (LLC-Nico) were maintained and treated similar to etomoxir-administrated mice, with the exception that 50 mg/kg body weight of nicotinic acid (Sigma-Aldrich, USA) dissolved in 5% (2-Hydroxypropyl)-β-cyclodextrin solution was i.p. injected daily from day 8 following tumor transplantation. Mice were sacrificed 19 days following tumor transplantation.

#### Glucose tolerance test

D-Glucose (2 g/kg body weight) was injected i.p. following a 6 h fast from 7:00 AM. Blood glucose levels before (0 minutes), 20 minutes, 40 minutes, 60 minutes and 90 minutes following glucose injection were measured using a glucometer (Bayer Contour, Switzerland). For measurements of plasma insulin concentration, blood was collected from the tail vein at time points 0 and 20 minutes. Plasma insulin was analyzed by ELISA in duplicates (Mouse Ultrasensitive Insulin ELISA, #80-INSTRU-E10, ALPCO Diagnostics, USA).

### 2.4. Body composition analysis

Fat mass and lean body mass were determined by quantitative magnetic resonance imaging (EchoMRI-4in1TM, Echo Medical System LLC, USA) 0-2 days before termination. Quantitative magnetic resonance imaging of the tumor itself disclose the tumor as 5% “fat mass” and 95% as “lean body mass” (unpublished observations), and therefore 5% and 95% of the dissected tumor mass was subtracted from the fat mass and lean body mass, respectively. *In vivo 2-deoxy-glucose uptake experiments.* To determine glucose uptake in skeletal muscle, perigonadal white adipose tissue (WAT), and the tumor,3H-labelled 2-deoxy-glucose ([3H]2DG) (Perkin Elmer, USA) was injected retro-orbitally (r.o.) in a bolus of saline (6 μl/g body weight) containing 66.7 μCi/ml [3H]2DG in mice as described [24]. The injectate also contained 0.3 U/kg body weight insulin (Actrapid; Novo Nordisk, DK) or a comparable volume of saline as well. Mice fasted for 3-5 h from 07:00 AM and were anesthetized (i.p. injection of 7.5 mg pentobarbital sodium per 100 g body weight) 15 minutes before the r.o. injection. Blood samples were collected from the tail vein and analyzed for glucose concentration using a glucometer (Bayer Contour, Switzerland). Sampling was performed immediately prior to insulin or saline injection and either after 5 and 10 minutes, or after 3, 6, 9, and 12 minutes as indicated in the figures. After 10 or 12 minutes, mice were humanely euthanized by cervical dislocation and the tumor, perigonadal WAT, tibialis anterior (TA), and gastrocnemius muscles were excised and quickly frozen in liquid nitrogen and stored at −80°C until processing. Once tissues were removed, blood was collected by punctuation of the heart, centrifuged (13,000 rpm, 5 minutes) and plasma frozen at −80°C. Plasma samples were analyzed for insulin concentration, IL-6, TNF-α, and specific [3H]2DG tracer activity. Plasma insulin was analyzed as described above. Tissue-specific 2DG uptake was analyzed as described [25, 26]. Plasma IL-6 and TNF-α were analyzed using V-PLEX Custom Mouse Cytokine Proinflammatory Panel1 (Mesoscale, #K152A0H-1) according to the manufacturer’s recommendation, loading 12.5 µl of plasma to each well.

Calculations of the whole-body glucose uptake index were performed as follows based on the body mass compositions obtained from the MRI scans: *y = a*b*, where *y* is 2DG uptake index (µmol/h), *a* is 2DG uptake rate (µmol/g/h), *b* is either tumor, fat, or 0.5*lean mass (g). Half of the lean mass was estimated to be muscle mass based on the study of Rolfe and Brown [27]. It was an assumption that all fat depots on average displayed glucose uptake similar to our measured WAT depot.

### 2.5. Microvascular perfusion in muscle

Mice were anesthetized with an i.p. injection of 11 μl/g body weight of Fentanyl (0.05 mg/ml, Dechra, DK), Midazolam (5 mg/ml, Accord Healthcare, UK) and Acepromazine (10 mg/ml, Pharmaxim, SE), and placed on a heating pad. In control (n=6) and large (tumor size > 800 mm^3^) tumor-bearing mice (n=6,), microvascular perfusion (MVP) was measured across the adductor magnus and semimembranosus muscles, with real-time contrast-enhanced ultrasound technique using a linear-array transducer connected to an ultrasound system (L9-3 transducer, iU22, Philips Ultrasound, Santa Ana, CA, USA) as described [28]. In short, a transducer was positioned over the left hindlimb and secured for the course of the experiment. A suspension of Optison microbubbles (Perflutren Protein-Type A Microspheres Injectable Suspension, USP, GE Healthcare, USA) was infused intravenously (15 μl/minute) using a Harvard 11 Plus low volume infusion pump (Harvard instrument Co., Holliston, MA). The infusion tube was attached to a vortex mixer to ensure a homogeneous microbubble solution entering the animal. An infusion time of 4 minutes was used where the first 2 minutes was to ensure systemic steady-state conditions before three consecutive MVP recordings were performed. Data were exported to quantification software (QLab, Philips, Andover, MA, USA) for analysis. Regions of interest were drawn clear of connective tissue and large vessels and copied into each file to ensure that regions were identical for each recording. Calculations were made in accordance with Wei et al. [29]. In short, acoustic intensity (AI) versus time curves were fitted to the exponential function: *y = A(1 – exp(−*β*(t − Bt))*, where *t* is time (seconds), *Bt* is the time used for background subtraction, *y* is the acoustic intensity at any given *t, A* is the plateau AI defined as MVP, and β is the flow rate constant (liters·s-1) that determines the rate of rising AI.

### 2.6. Basal and insulin-stimulated hepatic glucose production

Mice (control, n=5, or tumor-bearing mice (tumor size > 800 mm^3^), n=6) were clamped in randomized order after a 4 h fasting period from 10:00 AM. Mice were anesthetized with an i.p. injection of 11 μl/g body weight of Fentanyl (0.05 mg/ml, Dechra, DK), Midazolam (5 mg/ml, Accord Healthcare, UK) and Acepromazine (10 mg/ml, Pharmaxim, SE), and placed on a heating pad. A polyethylene cannula (PE50, Intramedic, USA) was inserted into a jugular vein for administration of anesthetics, insulin, and glucose. Anesthesia was maintained by constant infusion of the anesthetics (0.03 μl/g/min). After surgery, a 60 minutes continuous infusion (0.83 µl/minute, 1.2 µCi/h) of D-[3-3H]-glucose (Perkin Elmer) was administrated. Then, a 120 minutes hyperinsulinemic-euglycemic clamp was initiated, with a primed (4.5 mU) infusion of insulin (7.5 μU/kg/minute) (Actrapid, Novo Nordisk, DK) and D-[3-3H]-glucose (0.83 μl/minute, 1.2 μCi/h). Blood glucose was clamped at 6 mmol/l and maintained by a variable infusion of 20% glucose solution. Blood was sampled from the tail at −10, 0, 105, and 120 minutes for determination of plasma glucose, plasma 3H activity by scintillation counting, and thereby the plasma specific activity. Basal and insulin-stimulated HGP were calculated based on the equation described [30]. At 120 minutes, blood for plasma insulin concentration was also obtained from the tail. Mice were humanely euthanized by cervical dislocation.

### 2.7. Indirect calorimetry

After seven days of acclimation to single-housing, the mice were transferred to metabolic cages. Here, oxygen consumption and CO_2_ production were measured by indirect calorimetry in a CaloSys apparatus for 20 hours (TSE LabMaster V5.5.3, TSE Systems, GER) to calculate respiratory exchange ratio (RER). Non tumor-bearing mice were injected with a single dose of etomoxir (5 mg/kg body weight) or a vehicle control in a cross-over design with injections separated by 24 hours (n=6). All mice were acclimatized to thermoneutrality (30°C) for 3 weeks prior to the measurements.

### 2.8. Immunoblotting

Mouse muscles was pulverized in liquid nitrogen and homogenized 2 × 0.5 minutes at 30 Hz using a TissueLyser II bead mill (Qiagen, USA) in ice-cold homogenization buffer, pH 7.5 (10% glycerol, 1% NP-40, 20 mM sodium pyrophosphate, 150 mM NaCl, 50 mM HEPES (pH 7.5), 20 mM β-glycerophosphate, 10 mM NaF, 2 mM phenylmethylsulfonyl fluoride (PMSF), 1 mM EDTA (pH 8.0), 1 mM EGTA (pH 8.0), 2 mM Na_3_VO_4_, 10 μg/ml leupeptin, 10 μg/ml aprotinin, 3 mM benzamidine). After rotation end-over-end for 30 min at 4°C, supernatants from muscle tissue were collected by centrifugation (10,000 rpm) for 20 minutes at 4°C. Lysate protein concentrations were measured using the bicinchoninic acid method with bovine serum albumin (BSA) as a standard (Pierce). Total proteins and phosphorylation levels of relevant proteins were determined by standard immunoblotting techniques loading equal amounts of protein with a standard curve included for all proteins to ensure quantifications within the linear range. Polyvinylidene difluoride membranes (Immobilon Transfer Membrane; Millipore) were blocked in Tris-buffered saline (TBS)-Tween 20 containing 2% milk for 10-20 minutes at room temperature. Membranes were incubated with primary antibodies (Table 1) overnight at 4°C, followed by incubation with HRP-conjugated secondary antibody for 45 minutes at room temperature. Coomassie brilliant blue staining was used as a loading control [31]. Bands were visualized using the Bio-Rad ChemiDoc MP Imaging System and enhanced chemiluminescence (ECL+; Amersham Biosciences). Bands were quantified using Bio-Rad’s Image Lab software 6.0.1 (RRID:SCR_014210).

**Table 1.** Primary antibodies

| Antibody | Dilution (Primary) | Catalogue number | Company | RRID |
| --- | --- | --- | --- | --- |
| AKT2 | 1:1000 (in 2% milk) | #3063 | Cell Signaling Technology (CST) | AB_2225186 |
| P-AKT1/2<sup>Ser473/474</sup> | 1:1000 (in 2% milk) | #9271 | CST | AB_329825 |
| P-AKT1/2<sup>Thr308/309</sup> | 1:1000 (in 2% milk) | #9275 | CST | AB_329828 |
| TBC1D4 | 1:1500 (in 2% milk) | ab189890 | Abcam | – |
| P-TBC1D4<sup>Thr642</sup> | 1:1000 (in 2% milk) | #D27E6 | CST | AB_2651042 |
| Hexokinase II | 1:1000 (in 2% milk) | #2867 | CST | AB_2232946 |
| GLUT4 | 1:1000 (in 2% milk) | #PA1-1065 | Thermo Fisher Scientific | AB_2191454 |

### 2.9. RNA extraction and real-time PCR

RNA was isolated from 15 mg w.w. tumor tissue by guanidinium thiocyanate-phenol-chloroform extraction method with modifications [32], and tissue homogenized for 2 min at 30 Hz in a TissueLyser II (Qiagen, NL). cDNA was produced by reverse transcribing 3 µg of RNA using superscript II reverse transcriptase (Invitrogen, USA), and the samples diluted to 0.01 µg/ul. Content of interleukin 6 (IL-6), tumor necrosis factor α (TNF-α) and TATA box binding protein (TBP) were determined by real☐time PCR (ABI 7900 Prism, Applied Biosystems, US). Sequences used to amplify a fragment of IL-6 were FP: 5′GCTTAATTACACATGTTCTCTGGGAAA 3′, RP: 5′CAAGTGCATCATCGTTGTTCATAC 3′, Taqman probe: 5′ATCAGAATTGCCATTGCACAACTCTTTTCTCAT 3′, and for TNF-α FP: 5′ATGGCCCAGACCCTCACA 3′, RP: 5′TTGCTACGACGTGGGCTACA 3′, Taqman probe: 5′TCAGATCATCTTCTCAAAATTCGAGTGACAAGC 3′. TBP was used as housekeeping gene. The TBP probe was a pre-developed assay reagent from Applied Biosystems, US. TBP was similar between groups. All TaqMan probes were 5′-6-carboxyfluorescein (FAM) and 3′-6-carboxy-N,N,N′,N′-tetramethylrhodamine (TAMRA) labeled (Applied Biosystems, US) except TATA-binding protein (TBP) which was 5′ FAM with minor groove binding.

### 2.10. Statistical analyses

Results are shown as mean ± standard error of the mean (SE) with the individual values shown for bar graphs or mean ± SE for curve graphs. Statistical testing was performed using t-test, one-way or two-way (repeated measures when appropriate) ANOVA as applicable. Pearson’s correlation was used to test the relationship between the tumor volume and insulin-stimulated 2DG uptake or hepatic glucose production. Sidak post hoc test was performed for all ANOVAs to test the difference between LLC groups and the non-tumor control group. A log-transformation was performed in data-sets that were not normally distributed. Statistical analyses were performed using GraphPad Prism, version 7 (GraphPad Software, La Jolla, CA, USA, RRID: SCR_002798). For time-adjusted models of insulin-stimulated glucose uptake (related to Fig. 2) an adjusted model (Pearson’s correlations) was applied in IBM SPSS statistics 25 (IBM, USA, RRID: SCR_002865). The significance level for all tests was set at α = 0.05.

**Figure 1.**
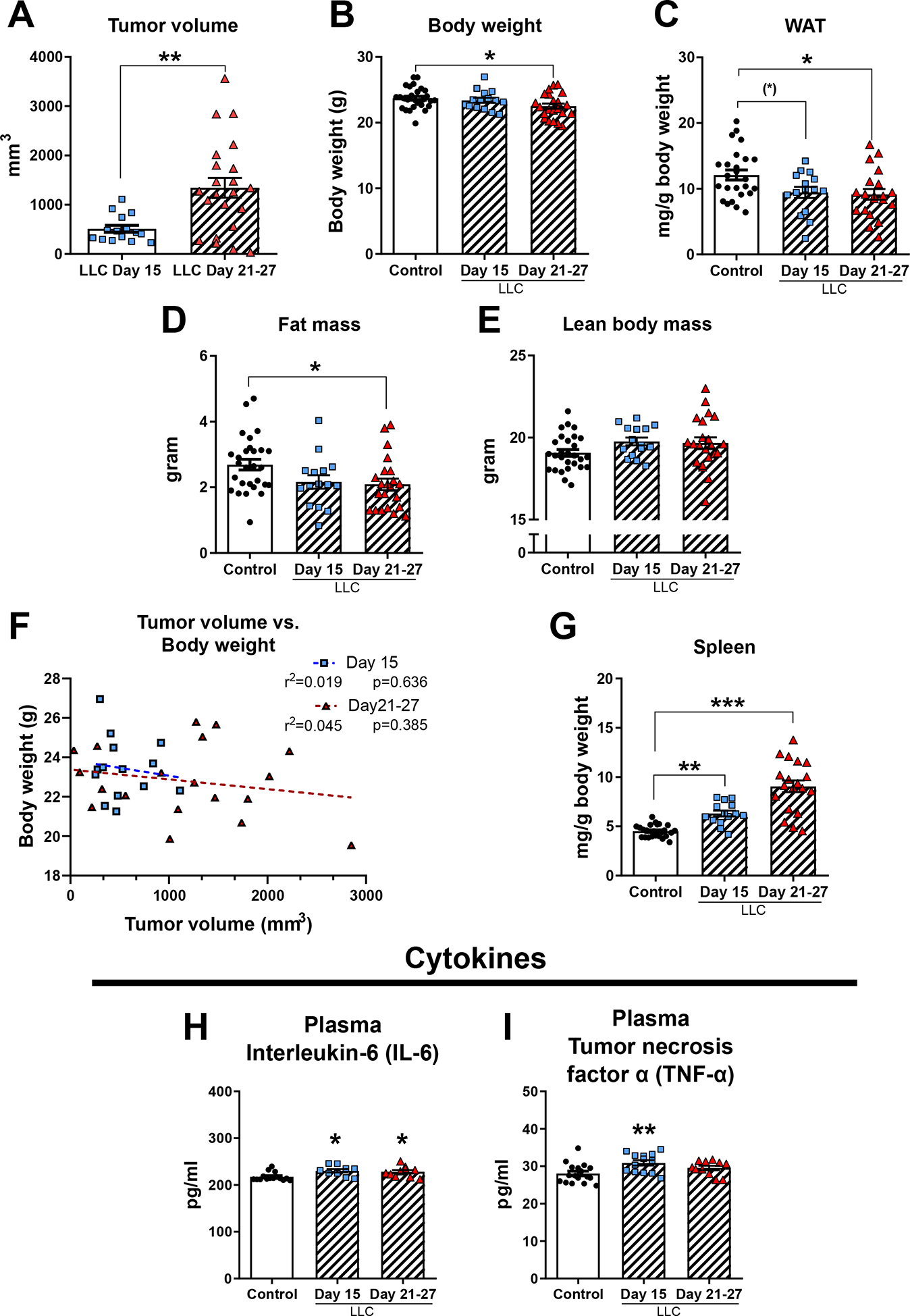
Characteristics of Lewis lung carcinoma (LLC) tumor-bearing mice. **A)** Tumor volume, **B)** body weight, **C)** perigonadal white adipose tissue (WAT) weight, **D)** fat mass, **E)** lean body mass, **F)** correlation between tumor volume and body weight, and **G)** spleen weight in control mice (n=25-28) and LLC tumor-bearing mice following 15 (n=14-15) or 21-27 days (n=19-22) tumor inoculation. The weights of all tissues were normalized to body weight with tumor weight subtracted. Plasma concentration of interleukin-6 (IL-6) (**H**) and tumor necrosis factor (TNF-α) (**I**) in control and tumor-bearing mice. Statistically significant effect of LLC on body composition or tissue weights is indicated by (*)P<0.1; *P < 0.05; **P < 0.01; ***P < 0.001. Values are shown as mean± SE with individual values.

**Figure 2.**
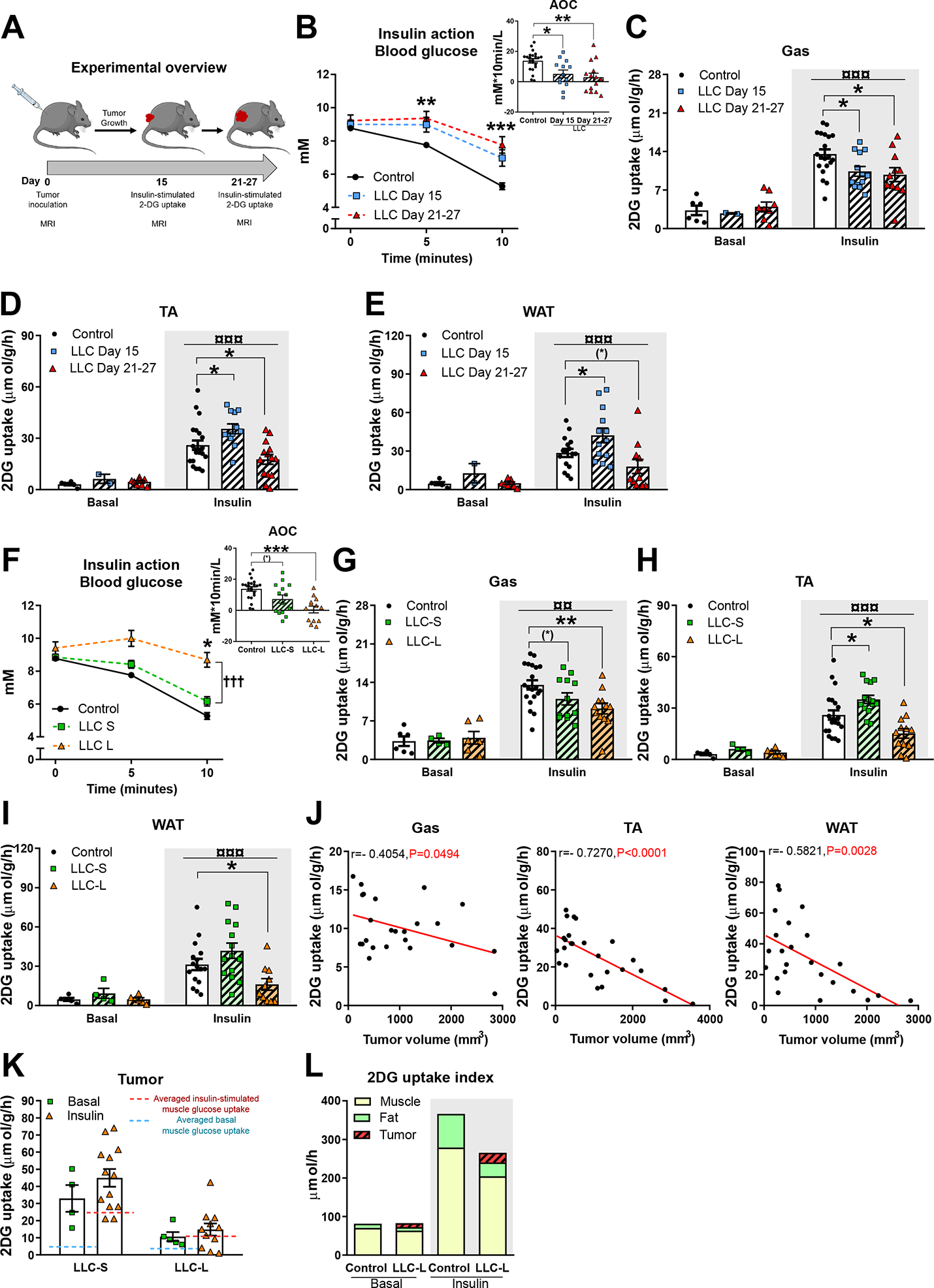
Insulin sensitivity in Lewis lung carcinoma (LLC) tumor-bearing mice. **A)** Experimental overview **B)** Blood glucose levels measured before (0 minutes), 5 minutes, and 10 minutes following retro-orbital (r.o.) insulin injection (0.3 U/kg body weight) and area over the curve (AOC) (n=22 in control group; n=13/14 in LLC groups) during the 10 minutes insulin stimulation. **C)** Basal (n=2-8) and insulin-(n=16-21 in control group; n=11-14 in LLC groups) stimulated 2-deoxy-glucose (2DG) uptake in gastrocnemius (Gas) muscle, **D)** tibialis anterior (TA) muscle, and **E)** perigonadal white adipose tissue (WAT) following 15 days or 21-27 days tumor inoculation. Data were re-divided into mice with large tumor volume (>800 mm^3^; LLC-L) and small tumors (<800 mm^3^; LLC-S). **F)** Blood glucose levels measured before (0 minutes), 5 minutes, and 10 minutes following r.o. insulin injection and AOC (n= 22 in control group; n= 13/14 in LLC groups) during the 10 minutes of insulin stimulation. **G)** Basal (n=4/6) and insulin (n= 16-21 in control group; n= 10-14 in LLC groups) stimulated 2DG uptake in Gas, **H)** TA, and **I)** WAT. **J)** Correlation between tumor volume and insulin-stimulated 2DG uptake in Gas, TA and WAT (n=24-26). **K)** Basal (n=4-5) and insulin-(n=12-14) stimulated glucose uptake in the tumor compared with muscle. **L)** Index of the contribution of muscle, fat, and tumor tissue to whole-body glucose uptake. Statistically significant effect of LLC on whole-body insulin action at each timepoint and 2DG uptake is indicated by (*)P<0.1; *P < 0.05; **P < 0.01; ***P < 0.001. Statistically significance on whole-body insulin action at 10 minutes between LLC-S and LLC-L is indicated by ††† P < 0.001. Statistically significant effect of insulin on 2DG uptake is indicated by ¤¤ P < 0.01; ¤¤¤P < 0.001.Values are shown as mean± SE with or without individual values.

## 3. Results

### 3.1. LLC caused adipose tissue wasting and increased spleen weight, indicative of mild cachexia and inflammation

Lewis lung carcinoma tumor-bearing day 15 mice had an average tumor volume of 530 mm^3^ ± 285 mm^3^. LLC day 21-27 mice had an average tumor volume of 1345 mm^3^ ± 946 mm^3^ (Fig. 1A). LLC day 21-27 tumor-bearing mice displayed reduced body weight (−5%; Fig. 1B), which was due to fat loss (Fig. 1C and D) rather than changes in lean body mass (Fig. 1E). Body weight was not associated with tumor-size (Fig. 1F). Spleen weight increased in LLC day 15 (+40%) and LLC day 21-27 (+101%) (Fig. 1G) tumor-bearing mice, indicating elevated immunomodulatory activity compared with control mice. This was confirmed by an average 7% increase in plasma levels of IL-6 (Fig. 1H) and TNF-α (Fig. 1I) in tumor-bearing mice. Thus, LLC induced adipose tissue wasting, increased spleen weight, and slightly elevated plasma cytokines, indicative of mild cachexia and inflammation.

### 3.2. LLC tumor-bearing mice displayed reduced blood glucose-lowering effect of insulin

We next investigated the effect of cancer on whole-body insulin action by retro-orbitally injecting a submaximal dose of insulin [33] and analyzing blood glucose in control and LLC tumor-bearing mice (Fig. 2A). In contrast to control mice, insulin did not lower blood glucose in LLC day 15 and LLC day 21-27 tumor-bearing mice 5 minutes following injection (Fig. 2B). Furthermore, the blood-glucose-lowering effect of insulin was markedly reduced (by 2-2.5 mM) at 10 minutes in LLC tumor-bearing mice compared to control mice (Fig. 2B). Accordingly, the area over the curve (AOC) was 60-80% lower in tumor-bearing mice (Fig. 2B), indicative of marked insulin resistance. Given the reduced whole-body insulin action, we measured glucose uptake in skeletal muscle and WAT, tissues that account for the majority of glucose utilization involved in whole-body glycemic control [34, 35]. In gastrocnemius muscle, insulin-stimulated glucose uptake was reduced in LLC day 15 tumor-bearing mice (−23%, Fig. 2C) and day 21-27 tumor-bearing mice (−28%, Fig. 2C) compared with control mice. Likewise, in TA muscle, insulin-stimulated glucose uptake was reduced in LLC day 21-27 tumor-bearing mice (−32%, Fig. 2D), but surprisingly, increased in LLC day 15 tumor-bearing mice (+45%, Fig. 2D). In perigonadal WAT, insulin-stimulated glucose uptake tended (P=0.0784) to be decreased by LLC at day 21-27 (−37%, Fig. 2E), while being increased in LLC at day 15 (+48%, Fig. 2E). Basal non-stimulated glucose uptake (Fig. 2C-E) and blood glucose (Fig. S1A) were unaffected by cancer. These findings show that cancer induces obvious insulin resistance and markedly reduces muscle and adipose tissue glucose uptake.

### 3.3. Insulin-stimulated glucose uptake in skeletal muscle and white adipose tissue negatively correlated with tumor size

As tissue-specific insulin resistance was more pronounced at day 21-27, we next investigated whether insulin resistance was related to the tumor size. We re-divided our current data set into small vs large tumors (cut off 800 mm^3^) across the LLC day 15 and LLC day 21-27 groups. In mice with large tumors (LLC-L), the blood-glucose-lowering effect of insulin was almost abrogated (Fig. 2F). However, in mice with small tumors (LLC-S), the blood glucose-lowering effect of insulin was only modestly reduced (Fig. 2F). Accordingly, AOC was 96% reduced in mice with large tumors and tended (P=0.051) to be 47% decreased in mice with small tumors (Fig. 2F).

In LLC-S tumor-bearing mice, insulin-stimulated glucose uptake tended (P=0.051) to be 19% decreased in gastrocnemius (Fig. 2G), increased in TA (+35%, Fig. 2H), while unaltered in WAT (Fig. 2I) compared to control mice. In contrast, LLC-L tumor-bearing mice displayed a marked reduction in insulin-stimulated glucose uptake in gastrocnemius (−32%, Fig. 2G), TA (−41%, Fig. 2H), and WAT (−44%, Fig. 2I). A negative correlation between tumor size and insulin-stimulated glucose uptake in both skeletal muscle and WAT (Fig. 2J) confirmed that cancer can significantly reduce insulin-stimulated glucose uptake in skeletal muscle and WAT in a tumor size-dependent manner. As the size of the tumor increases with time, we adjusted the above model for the duration of tumor-bearing. The oberserved negative correlation between tumor size and insulin-stimulated glucose uptake persisted when taking tumor burden duration into account (Supplementary Table 1). We observed no correlations between plasma IL-6 (Fig. S1B) or TNF-α (Fig. S1C) and insulin-stimulated glucose uptake in skeletal muscle or WAT. In the tumor tissue the mRNA expression of IL-6 and TNF-α was 60% and 20% higher, respectively, in day 15 compared to day 21-27 tumor-bearing mice (Fig. S1D). Surprisingly, we observed a positive correlation between tumor IL-6 (Fig. S1E) and TNF-α (Fig. S1F) expression and insulin-stimulated glucose uptake in TA and WAT. Nonetheless, with no correlation between plasma cytokines and glucose uptake and a positive correlation with the expression of tumor cytokines, systemic inflammation is likely not a main driver of insulin resistance in tumor-bearing mice in the present study.

Previous studies have reported that tumors take up a substantial amount of glucose to support tumor growth, migration, and invasion [36, 37]. However, to the best of our knowledge, no study has to date compared glucose uptake into muscle and tumor in the same mouse. We, therefore, analyzed tumor glucose uptake and found that tumor glucose uptake per gram tumor mass was 5.9- and 1.7-fold higher in LLC-S and LLC-L, respectively compared with average basal muscle glucose uptake (Fig. 2K, P<0.05). Similar results were obtained in the insulin-stimulated state, where the average muscle glucose uptake was 40% that of glucose uptake in tumors of LLC-S tumor-bearing mice (p< 0.01). Insulin-stimulated glucose uptake was similar in the tumor and muscle of LLC-L tumor-bearing mice (Fig. 2K). Those findings suggest that the tumor significantly competes with skeletal muscle for glucose. In order to understand how much the tumor contributed to whole-body glucose disposal during the 10 minutes of stimulations, we produced an index of whole-body glucose disposal in fat, skeletal muscle, and the tumor. It showed that in cancer, a substantial part of the available glucose is directed from muscle and fat into the tumor (Fig. 2L).

### 3.4. Reduced skeletal muscle glucose uptake in tumor-bearing mice was not due to impaired AKT signaling

Accounting for the majority of insulin-stimulated glucose disposal [35], we next determined canonical insulin signaling in skeletal muscle to elucidate the molecular mechanisms underlying LLC-induced insulin resistance. We focused our investigation on mice with tumor size of more than 800 mm^3^ (average tumor volume 1755 ± 763 mm^3^) as insulin resistance was pronounced in this group. To our surprise, we found upregulated phosphorylation of AKT^Ser474^ (+65%, Fig. 3A), AKT^Thr309^ (+86%) (Fig. 3B), and TBC1D4^Thr642^ (+18%, Fig. 3C), indicative of enhanced insulin sensitivity. Protein expression of AKT2 (Fig. 3D), TBC1D4 (Fig. 3E), glucose transporter 4 (GLUT4) (Fig. 3F), and Hexokinase II (Fig. 3G) remained unaltered in tumor-bearing mice, although we did detect a tendency (p=0.053) to increased protein expression for TBC1D4 in tumor-bearing mice (representative blots are shown in in Fig. 3H-I).

**Figure 3.**
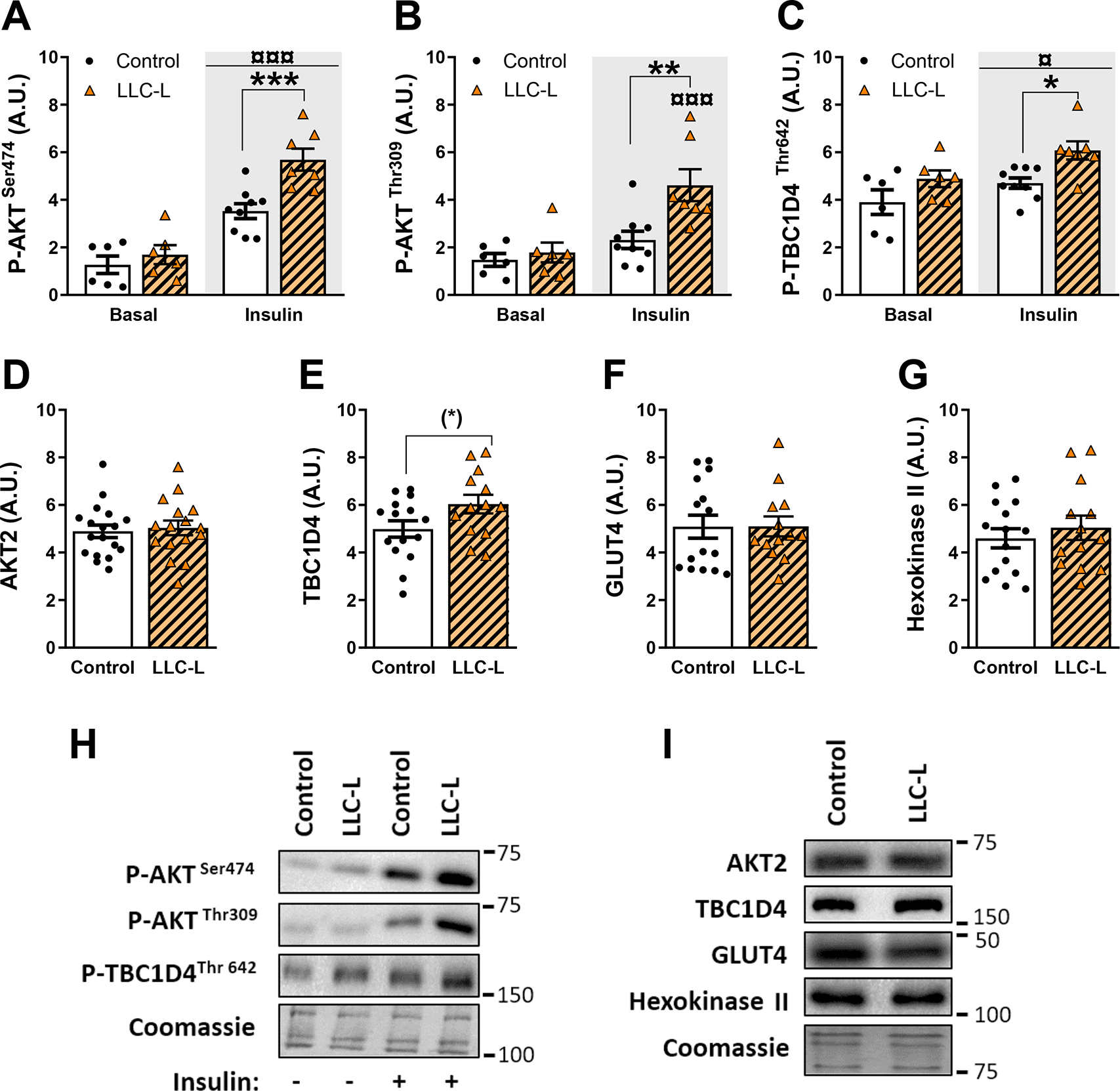
Effect of Lewis lung carcinoma on insulin-stimulated signaling in gastrocnemius muscle of mice with large tumors (>800mm^3^; LLC-L). **A)** Basal (n=6) and insulin-(n=10-12) stimulated phosphorylated (P)-AKT^Ser474^, **B)** P-AKT^Thr309^, **C)** P-TBC1D4^Thr642^. **D)** Protein expression of AKT2, **E)** TBC1D4, **F)** GLUT4, and **G)** Hexokinases II (n=16-18). **H)** Representative phospho-blots. **I)** Representative blots of total proteins. Statistically significant effect of LLC-L on insulin signaling is indicated by (*)P<0,1; *P < 0.05; **P < 0.01; ***P < 0.001. Statistically significant effect of insulin is indicated by ¤ P < 0.05; ¤¤¤P < 0.001.Values are shown as mean ± SE with individual values. A.U., arbitrary units.

These findings suggest that reduced insulin-stimulated muscle glucose uptake in tumor-bearing mice is not due to decreased myocellular canonical insulin signaling, on the contrary, we observed augmented phosphorylation of both AKT and TBC1D4.

### 3.5. Cancer abrogated muscle microvascular perfusion in response to insulin

Muscle microvascular perfusion (MVP) is essential for insulin to fully stimulate glucose uptake in muscle, however, it is unknown whether cancer influences muscle MVP. Thus, we determined muscle MVP in mice with tumor sizes > 800 mm^3^ (LLC-L, averaged tumor volume 1283 ± 207 mm^3^). In control mice, insulin increased muscle MVP (+70%; Fig. 4A and B) in accordance with previous studies [38, 39]. Remarkably, this increase was completely abrogated in LLC-L tumor-bearing mice (Fig. 4C and D), showing that cancer negatively affects insulin-stimulated muscle microvascular perfusion, which could contribute to muscle insulin resistance.

**Figure 4.**
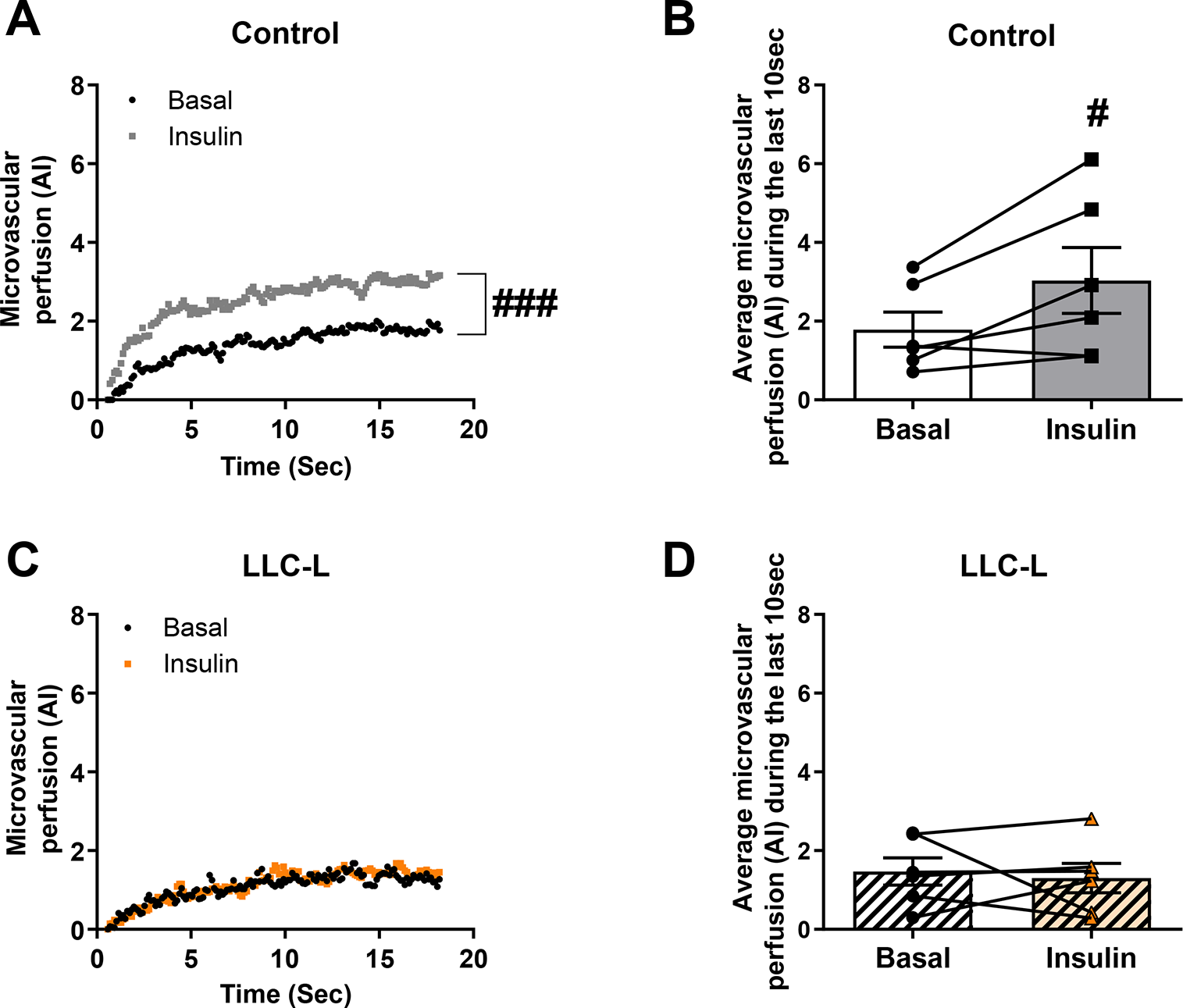
Effect of Lewis lung carcinoma on skeletal muscle microvascular perfusion in mice with large tumors (>800mm^3^; LLC-L). **A)** Microvascular refilling curves after microbubbles destruction **B)** microvascular perfusion presented as the plateau AI value in adductor magnus and semimembranosus muscles in control (**A** and **B**) and LLC-L (**C** and **D**) tumor-bearing mice at baseline and after 60 minutes of insulin (7.5 μU /kg/minute) infusion (n=6). Statistically significant effect of insulin on microvascular perfusion is indicated by #P < 0.05; ###P < 0.001. Values are shown as mean or mean ± SE with individual values.

### 3.6. Tumor-bearing mice exhibit increased basal hepatic glucose production

Increased hepatic glucose production (HGP) is another hallmark of insulin resistance. Therefore, we measured basal and insulin-stimulated HGP in mice with tumor sizes > 800 mm^3^ (LLC-L, averaged tumor volume 3916 ± 2196 mm^3^) during a hyperinsulinemic-euglycemic clamp. Following 120 minutes of continuous insulin infusion (7.5 μU/kg/minute), blood glucose in both control and LLC-μL tumor-bearing mice was maintained at a steady level of 6 mM (Fig. 5A). Steady-state glucose infusion rate (GIR) during the clamp was similar between control and tumor-bearing mice (Fig. 5B). This is in contrast to our findings during the 10 minutes insulin stimulation where tumor-bearing mice displayed reduced insulin response. This discrepancy might reflect the fact that the tumor takes up a large proportion of the glucose in the tumor-bearing mice as indicated in Fig. 2K and L. Over time that could masks smaller reductions in muscle and adipose tissue glucose uptake. It could also reflect the longer stimulation time (the last 30 minutes of the clamp used to calculate GIR), or that the insulin dose used to estimate GIR was higher compared with the r.o. injection.

**Figure 5.**
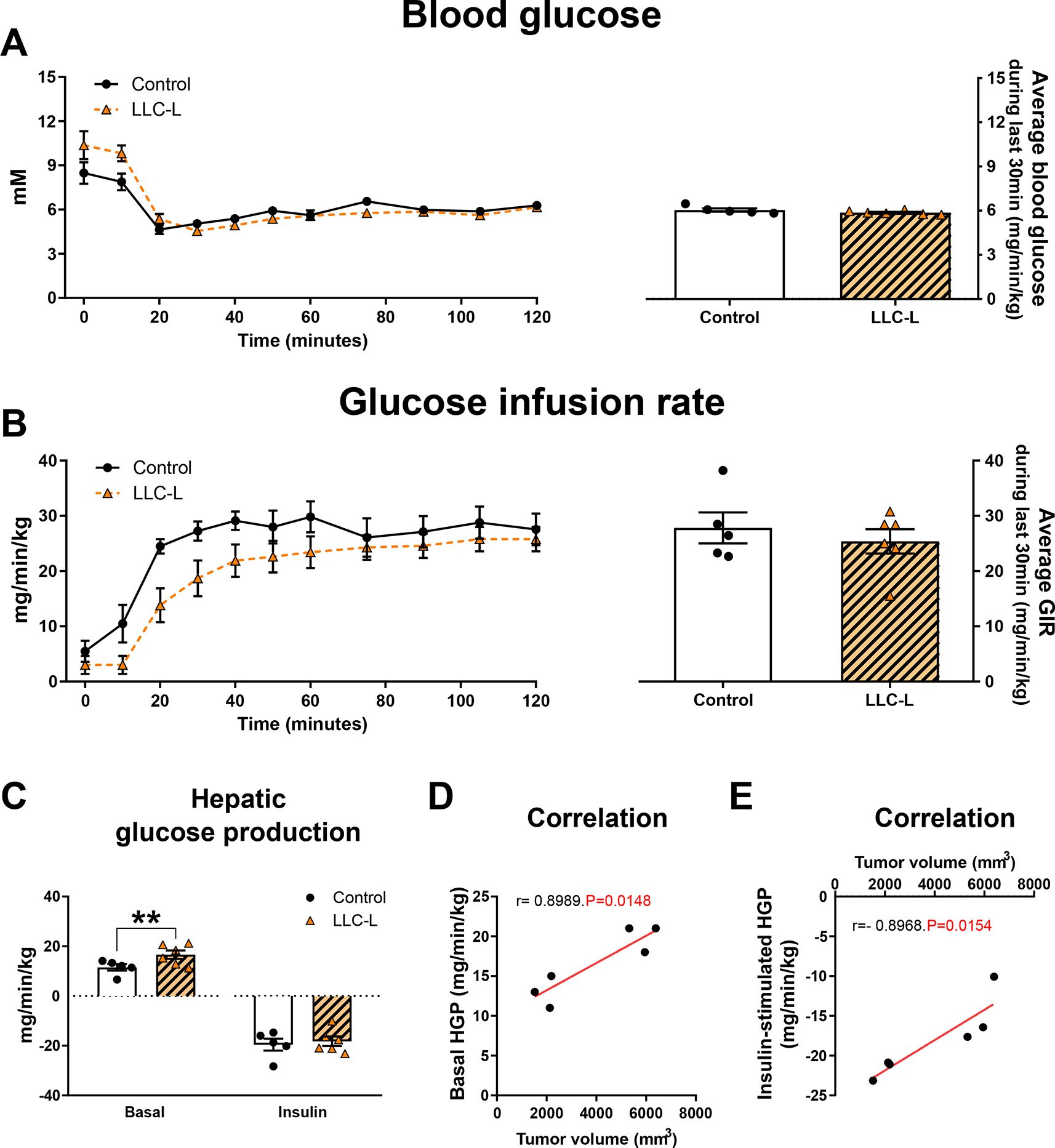
Effect of Lewis lung carcinoma on hepatic glucose production (HGP) in mice with large tumors (>800mm^3^; LLC-L). **A)** Blood glucose, **B)** glucose infusion rate (GIR), **C)** basal or insulin-stimulated HGP in control mice (n=5) and LLC-L tumor-bearing mice (n=6), **D)** correlation between tumor volume and basal or **E)** insulin-stimulated HGP during hyperinsulinemic-euglycemic clamp (7.5 μU /kg/minute). Statistically significant effect of LLC-L on basal HGP is indicated by **P < 0.01. Values are shown as mean ± SE with or without individual values.

Interestingly, basal HGP was 45% higher in LLC-L tumor-bearing mice compared to controls (Fig. 5C). In addition, basal HGP positively correlated with tumor volume (Fig. 5D). Insulin suppressed HGP similarly in LLC-L tumor-bearing and control mice (Fig. 5C). Nevertheless, within the tumor-bearing group, the inhibitory effect of insulin on HGP was negatively correlated with tumor volume (Fig. 5E), although this should be interpreted cautiously, given the low number of mice. Collectively, these findings show that cancer increases basal HGP, but does not affect insulin-stimulated GIR or insulin-suppressed HGP at supra-physiological insulin levels.

### 3.7. Inhibition of fatty acid oxidation partially restored LLC-induced insulin resistance and inhibition of lipolysis improved glucose tolerance

Augmented fatty acid metabolism, a hallmark of many insulin resistance conditions [16,40,41], has been reported in human [42] and murine [19, 20] cancer models. Therefore, we tested the hypothesis that cancer might reduce insulin action via its effect on fatty acid metabolism in LLC tumor-bearing mice (averaged tumor volume 645 ± 386 mm^3^). To inhibit fatty acid oxidation, we used the CPT1 inhibitor, etomoxir. In non-tumor-bearing mice, etomoxir treatment increased plasma free fatty acids (Fig. 6A) and acutely increased RER (Fig. 6B) as would be expected by inhibition of fatty acid oxidation [43]. In agreement with previous reports in cancer patients [13–15] and mouse cancer models [19, 20], we observed increased plasma free fatty acids (FFA) (+125%, Fig. 6C), triacylglycerol (+23%, Fig. 6D) and glycerol (+40%, Fig. 6E) concentrations in LLC tumor-bearing compared to control mice.Daily administration of etomoxir, showed that inhibition of whole-body fatty acid oxidation restored blood free fatty acid concentrations to levels of control mice (Fig. 6C-E). Plasma IL-6 (+17%, Fig. 6F) and TNF-α (p=0.09, +10%, Fig. 6G) were increased in tumor-bearing mice. In contrast to the plasma lipid profile, etomoxir treatment did not affect the plasma levels of the cytokines IL-6 and TNF-α (Fig. 6F-G). Similar spleen weight was observed tumor-bearing mice with or witout etomoxir treatment (Fig. S2A). Thus, inhibition of fatty acid oxidation, at least partially, corrects abnormal lipid metabolism induced by cancer in mice as measured by plasma triacylglycerol, free fatty acids, and glycerol.

**Figure 6.**
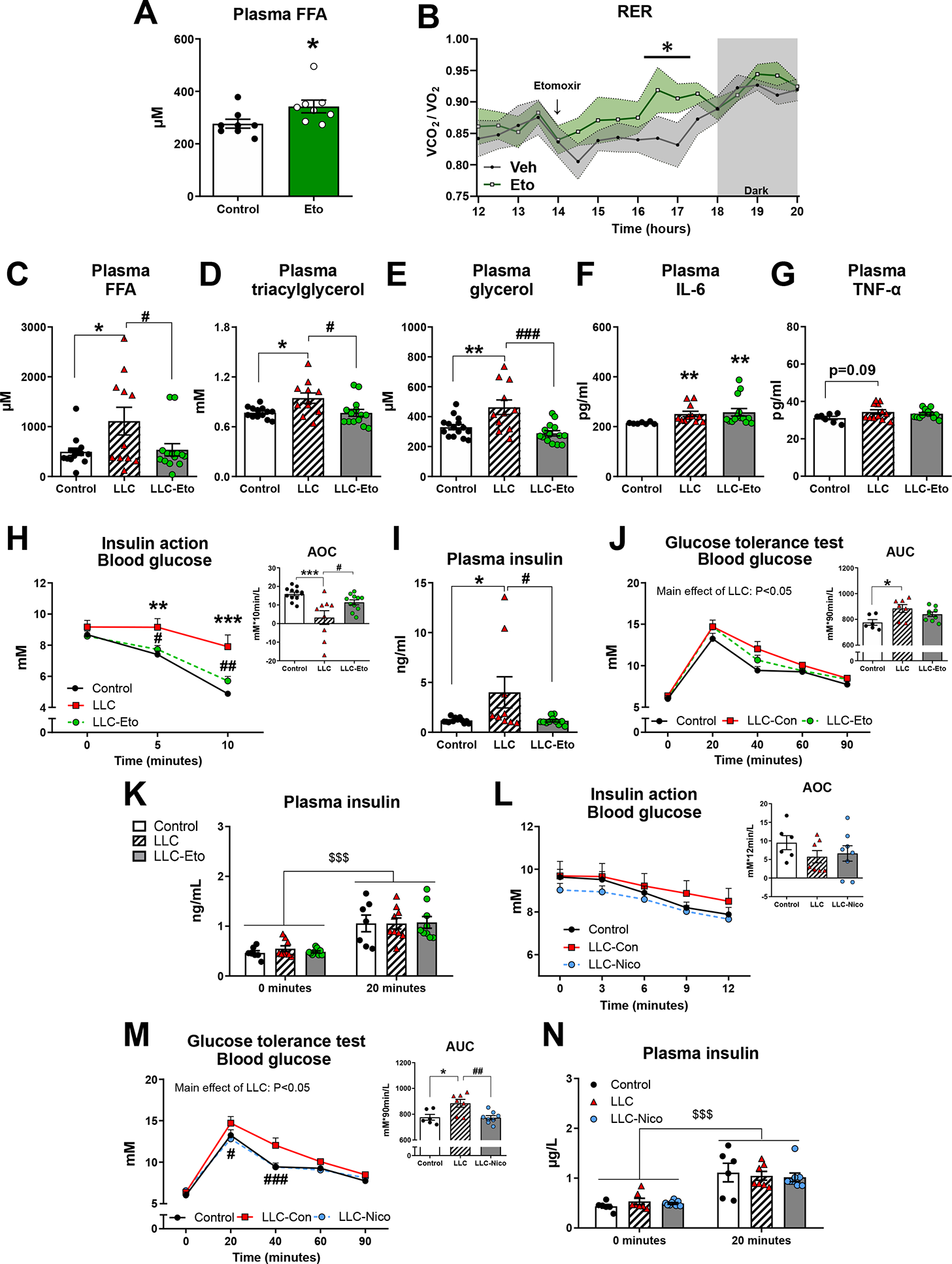
Effect of etomoxir on insulin sensitivity and glucose tolerance. **A)** acute effect of etomoxir treatment in non-tumor-bearing mice (n=8). **B)** acute effect of etomoxir treatment on respiratory exchange ration (RER) (n=6). **C)** Plasma free fatty acid (FFA), **D)** triacylglycerol, **E)** glycerol, **F)** plasma interleukin-6 (IL-6), and **G)** plasma tumor necrosis factor α (TNF-α) in control mice and LLC tumor-bearing mice with or without etomoxir (Eto) administration (n=11-15). **H)** Blood glucose levels before (0 minutes), 5 minutes and 10 minutes following retro-orbital insulin injection (0.3 U/kg body weight) and area over the curve (AOC). **I)** Plasma insulin (n=9-12). **J)** Blood glucose levels before (0 minutes), 20 minutes, 40 minutes, 60 minutes and 90 minutes following an intraperitoneal glucose tolerance test (2 g/kg body weight) and area under the curve (AUC) (n=6-9). **K)** Plasma insulin levels at 0 minutes and 20 minutes following intraperitoneal glucose tolerance test (2 g/kg body weight) (n=6-9). **L)** Blood glucose concentration before (0 minutes), 5 minutes and 10 minutes following retro-orbital insulin injection (0.3 U/kg body weight) in control mice or Lewis lung carcinoma (LLC) tumor-bearing mice with or without nicotinic acid (Nico) administration (n=8-12), and area over the curve (AOC). **M)** Blood glucose concentration before (0 minutes), 20 minutes, 40 minutes, 60 minutes and 90 minutes following intraperitoneal glucose tolerance test (GTT; 2 g kg-1 body weight) in control and LLC tumor-bearing mice with or without nicotinic acid (Nico) administration and area under the curve (AUC). **N)** plasma insulin concentration at 0 minutes and 20 minutes into the GTT (n=6-8). Statistically significant effect of LLC is indicated by *P < 0.05; **P < 0.01; ***P < 0.001. Statistically significant effect of Eto/Nico is indicated by #P < 0.05; ##P < 0.01; ###P < 0.001. Values are shown as mean± SE with or without individual values.

Next, we investigated whether the inhibition of fatty acid oxidation improved the metabolic dysfunction of tumor-bearing mice. In control mice, insulin lowered blood glucose by 2 mM and 4 mM following 5 and 10 minutes of stimulation, respectively (Fig. 6H). In agreement with our previous observations in the present study, insulin action in LLC tumor-bearing mice was reduced (Fig. 6H). Remarkably, etomoxir rescued insulin action in LLC tumor-bearing mice evidenced by restored blood glucose levels (Fig. 6H) and 2.5-fold increase of AOC (Fig. 6H). We also analyzed plasma insulin concentration 10 minutes following the r.o. insulin injection, as a marker of insulin clearance. Interestingly, LLC tumor-bearing mice showed increased plasma insulin (+237%) (Fig. 6I), an indication of reduced insulin clearance, which is also observed in diet-induced insulin-resistant mice [24, 44]. Etomoxir administration normalized plasma insulin levels in LLC tumor-bearing mice (Fig. 6I). In order to evaluate glycemic regulation, we undertook a glucose tolerance test and found that tumor-bearing mice were glucose intolerant (Fig. 6J). Glucose intolerance was not rescued by etomoxir (Fig. 6J), despite the improvements in circulating fatty acid levels and insulin action. The glucose challenge increased plasma insulin levels 100-150% similarly in all groups (Fig. 6K). Tumor volume was not affected by etomoxir treatment (Fig. S2B) and food intake was similar between all groups (Fig. S2C). Etomoxir did not affect insulin’s blood glucose lowering effect in non-tumor-bearing mice (Fig. S2D), suggesting that the benefits of inhibiting fatty acid oxidation on insulin sensitivity were specific to the context of cancer-induced accelerated fatty acid metabolism.

We next inhibited adipose tissue lipolysis by a potent inhibitor, nicotinic acid. Nicotinic acid did not increase insulin’s blood glucose-lowering effect in non-tumor-bearing mice (Fig. S2E), nor did it affect glucose tolerance (Fig. S2F). Tumor size and growth rate were not affected by nicotinic acid treatment (Fig. S2G) and spleen weight increased similarly in tumor-bearing mice with or without nicotinic acid treatment (Fig. S2H). In contrast to etomoxir, the blood glucose response to r.o. injected insulin was not improved by nicotinic acid (Fig. 6L). However, nicotinic acid did restore glucose tolerance in LLC-tumor-bearing mice (Fig. 6M). The glucose challenge increased plasma insulin levels similarly (100-150%) in all groups (Fig. 6N), suggesting that altered insulin sensitivity rather than insulin levels caused the improvement in glucose tolerance by nicotinic acid. Taken together, these findings demonstrate that altered fatty acid metabolism, but likely not inflammation, is involved in LLC-induced insulin resistance and glucose intolerance.

## 4. Discussion

Cancer resulted in marked perturbations on at least six metabolically essential functions; i) insulin’s blood-glucose-lowering effect, ii) glucose tolerance, iii) skeletal muscle and WAT insulin-stimulated glucose uptake, iv) intramyocellular insulin signaling, v) muscle microvascular perfusion, and vi) basal hepatic glucose production in mice (depicted in Fig. 7). Additionally, we show that the mechanism causing cancer-induced insulin resistance may relate to altered fatty acid metabolism but is likely not related to inflammation.

**Figure 7.**
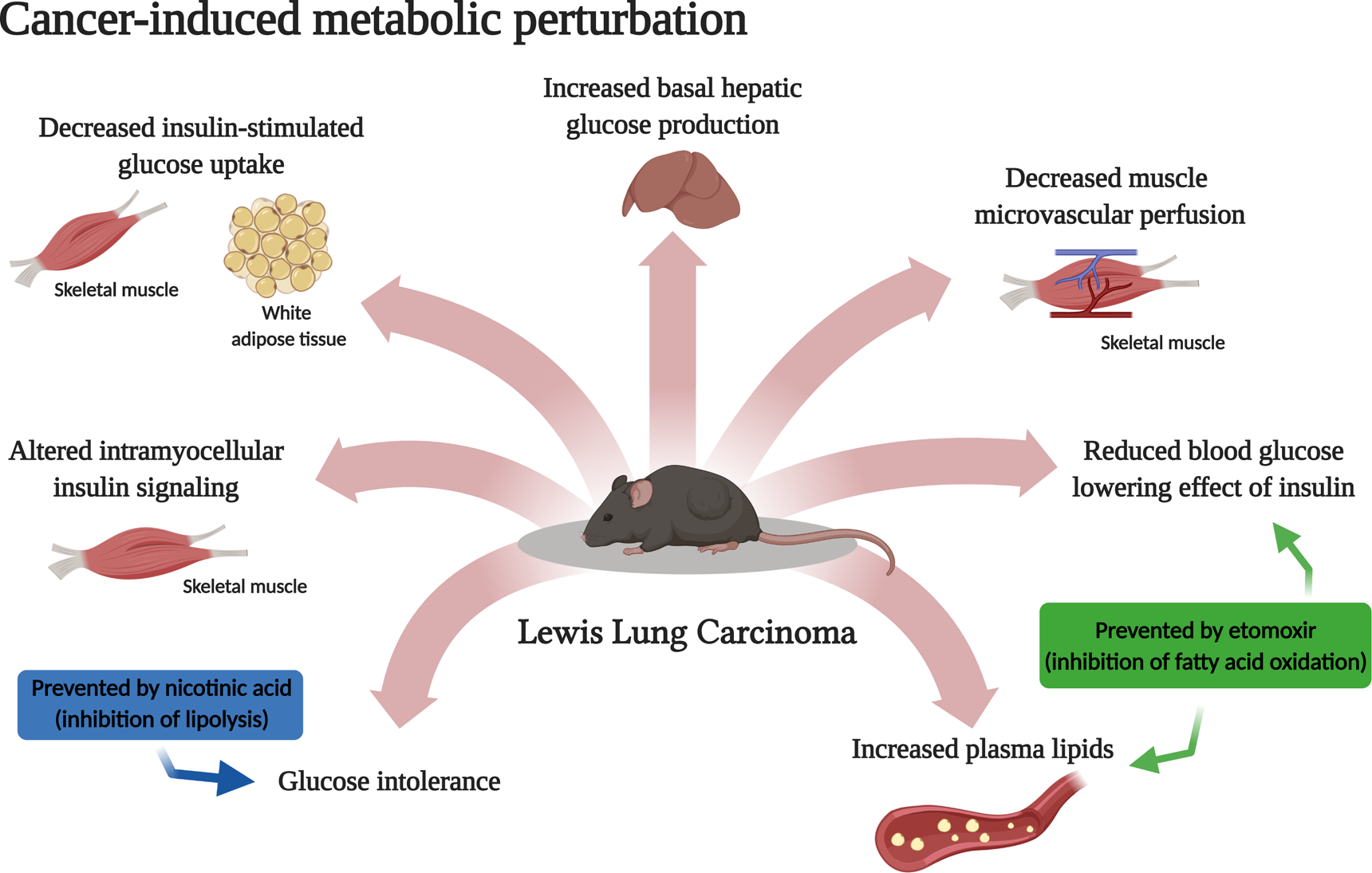
Graphic overview of Lewis lung cancer-induced metabolic perturbations identified by the present study. Created with BioRender.com.

A major finding in the current study was the significantly impaired insulin-stimulated glucose uptake in both skeletal muscle and adipose tissue in LLC tumor-bearing mice. These findings suggest that skeletal muscle and adipose tissue are major players in dysregulated glucose metabolism often observed in human cancers and murine cancer models [45–47]. In accordance, reduced insulin-stimulated glucose disposal has also been reported in cachexic patients with lung cancer [8] and lymphoma [48], suggesting that our findings are clinically relevant and translatable to the human situation. Tumor size seemed to be a key factor in peripheral insulin resistance, as we observed the skeletal muscle and WAT glucose uptake to be negatively correlated with tumor size. Conversely, inflammation seemed not to be involved in cancer-associated insulin resistance, as plasma IL-6 and TNF-α did not correlate with glucose uptake rates. Interestingly, the decreased insulin-stimulated glucose uptake in mice with large tumors was not due to diminished proximal insulin signaling in muscle. On the contrary, insulin-stimulated AKT/TBC1D4 signaling was upregulated in skeletal muscle of tumor-bearing mice. This is surprising, given that tumor-bearing mice displayed increased whole-body inflammation as indicated by increased spleen volume and increased plasma levels of IL-6 and TNF-α, consistent with other investigations showing increased IL-6 in patients with non-small cell lung cancers [49, 50]. Inflammation would be expected to reduce insulin signaling in muscle [51]. Tumorkines, such as VEGF and HIF-1, are reported to be upregulated in the LLC model of cancer [52, 53]. Although not analyzed in our study, those tumorkines could increase insulin signaling in muscle, as they have been reported to increase PI3K/AKT signaling in cancer cells [37, 54]. The causes of upregulated muscle insulin signaling in cancer warrants further investigation but increased insulin signaling in muscles with reduced insulin-stimulated glucose uptake has been reported in other models [55]. Nevertheless, the mechanisms by which cancer causes insulin resistance seems to be different from the mechanisms causing insulin resistance in for example obesity and type 2 diabetes, where muscle AKT and TBC1D4 signaling is either unaffected [56, 57] or reduced [58–60]. Furthermore, inflammation did not seem to be a main driver of insulin resistance, as spleen volume, plasma IL-6, and plasma TNF-α were similarly increased in mice with small and large tumors, of which only the mice with large tumors displayed reduced insulin-stimulated glucose uptake in muscle and WAT.

Another major finding of the present investigation was that insulin-stimulated muscle microvascular perfusion was abrogated in tumor-bearing mice. To our knowledge, these data are the first to show that dysregulated muscle microvascular perfusion is involved in a common model of cancer and cachexia. Insulin-stimulated microvascular perfusion in muscle is a critical facet in glucose uptake regulation [10,39,61–63]. In agreement, genetic or pharmacological inhibition of microvascular perfusion impaired insulin-stimulated muscle glucose uptake by 40% in otherwise healthy mice [38]. In addition, insulin-stimulated microvascular perfusion is reduced in different insulin-resistant conditions, including obesity and type 2 diabetes [64, 65]. In cancer, impaired microvascular perfusion could be caused by elevated circulating fatty acid levels, as experimentally elevated circulating fatty acids reduced insulin-stimulated muscle microvascular perfusion by 40% [66, 67] without causing impairments in intracellular insulin signaling in healthy humans [17]. Our findings show that decreased microvascular perfusion could contribute to cancer-induced impaired muscle glucose uptake in response to insulin.

In our study, a reduction in skeletal muscle and WAT glucose uptake likely contributed to the attenuated blood-glucose-lowering effect of insulin in tumor-bearing mice. On the other hand, Lang et al [68] have previously reported that insulin resistance in tumor-bearing rats was due to an impaired ability of insulin to suppress hepatic glucose production, although that study did not analyze muscle and adipose tissue glucose uptake. In the present study, insulin’s inhibitory effect on hepatic glucose production was not impaired, while basal hepatic glucose production was 45% increased. Increased basal hepatic glucose production has also been reported in patients with lung cancer [69] but the mecahnisms remain to be established.

Elevation of fatty acids is often associated with insulin resistance [16–18] and increased plasma free fatty acid concentrations are reported in cancers [19,20,46,70]. For example, the release of fatty acids and glycerol from WAT explants was 30-40% increased in LLC or B16 tumor-bearing mice [19]. Increased circulating fatty acids have been shown to induce insulin resistance [71]. Indeed, we found that whole-body insulin action was restored by blocking fatty acid oxidation via etomoxir administration in tumor-bearing mice. Furthermore, lipolysis inhibition via nicotinic acid administration rescued glucose intolerance in tumor-bearing mice. Interestingly, etomoxir normalized plasma triacylglycerol, fatty acids, and glycerol concentrations in tumor-bearing mice, which could benefit insulin action. The plasma fatty-acid-lowering effect of etomoxir in tumor-bearing mice is somewhat surprising, as fatty acid oxidation inhibition in non-tumor-bearing mice increased circulating fatty acid levels in our present and other studies [72, 73], and prolonged FA oxidation inhibition thus increased plasma triacylglycerol and liver fat content [43]. However, etomoxir inhibits lipolysis in adipocytes and can increase re-esterificationof fatty acids to triacylglycerol in the liver, thereby diminishing the release of fatty acids from the adipose tissue and triacylglycerol from the liver [74]. In the cancer condition with accelerated lipolysis [13, 15], it is possible that etomoxir’s effect on triacylglycerol metabolism seen in liver and lipolysis overrules the contrary effect of etomoxir on reduced fatty acid oxidation. Etomoxir has also been reported to reduce inflammation [75, 76], which could also contribute to the amelioration of cancer-induced insulin resistance. However, spleen weight and plasma IL-6 and TNF-α levels increased similarly in tumor-bearing mice with or without etomoxir treatment, suggesting that etomoxir did not prevent cancer-induced inflammation. Indeed, our results indicate that the impact of cancer on lipid metabolism exerts a critical impact on the pathology of cancer-induced insulin resistance.

Based on the present investigation and relevant literature [13–18,77], we hypothesize a model where the tumor secretes tumorkines that increase fatty acid metabolism, which in turn leads to peripheral insulin resistance. Redirecting glucose from skeletal muscle and adipose tissue, likely benefits the tumor’s energy demand to support tumor growth, migration, and invasion [36]. The clinical relevance of this is suggested, as cancer patients with type 2 diabetes have increased mortality rates [2].

There are certain limitations to the present investigation and unresolved questions. Firstly, our investigation was undertaken in a mouse model of lung cancer with a subcutaneously implanted tumor. Whether our results apply to the human condition and other types of cancer remain to be determined. However, patients with lung cancer have been reported to display reduced insulin-stimulated glucose uptake [8] and accelerated hepatic glucose production [69], suggesting that our findings do mimic the human condition. Secondly, insulin resistance was observed without loss of body mass. While this provides a good model for investigating the effect of cancer isolated from the presence of cachexia, we cannot draw any conclusions regarding the involvement of insulin resistance in cancer cachexia or *vice versa* [4–6]. Thirdly, our study shows that MVP was markedly compromised by cancer. However, whether this was due to altered fatty acid metabolism remains undetermined and future studies should investigate the effect of etomoxir and nicotinic acid on MVP. Treatment with pharmacological vasodilators have shown similar [78] or increased [79] muscle glucose uptake, thus, whether vasodilators could improve insulin sensitivity in cancer is also an unresolved question to be answered by future investigations.

In conclusion, cancer impaired the blood-glucose-lowering effect of insulin, caused glucose intolerance, and reduced glucose uptake in muscle and WAT. Furthermore, tumor-bearing mice displayed increased basal hepatic glucose production. Cancer-associated insulin resistance was neither due to inflammation nor impaired proximal muscle insulin signaling, but was associated with a complete abrogation of insulin-stimulated muscle microvascular perfusion. Finally, we identify fatty acid metabolism as a player in cancer-associated insulin resistance, providing potential therapeutic targets for cancer-induced insulin resistance. These findings suggest that insulin resistance is likely of key importance in the therapy of cancer.

## Author contributions

Author contributions. X.H., S.H.R., and L.S. conceptualized and designed the study. X.H., S.H.R., M.C., and L.S. conducted the experiments, performed the laboratory analysis, analyzed the data. X.H., S.H.R., and L.S. wrote the manuscript. K.A.S., C.H-O., M.A., A.M.L., A.M.F.., L.L.V.M., Z.L., J.L.,T.E.J., and B.K. all took part in conducting the experiments, performing laboratory analysis and/or interpreting the data. All authors commented on and approved the final version of the manuscript. All authors are guarantor of this work and take responsibility for the integrity of the data and the accuracy of the data analysis.

## Acknowledgments

We acknowledge the highly skilled technical assistance of Betina Bolmgren and Irene Bech Nielsen (Molecular Physiology Group, Department of Nutrition, Exercise and Sports, Faculty of Science, University of Copenhagen, Denmark). We also acknowledge the methodological assistance and loan of qPCR equipment by Professor Henriette Pilegaard (Section for Cell Biology and Physiology, Department of Biology, University of Copenhagen, Denmark). Finally, we wish to express our sincere gratitude and acknowledge Pernille Højman (Centre of Inflammation and Metabolism, Rigshospitalet, Copenhagen, Denmark) for invaluable discussions and methodological insights on this study.

---

**Figure supplementary 1.**
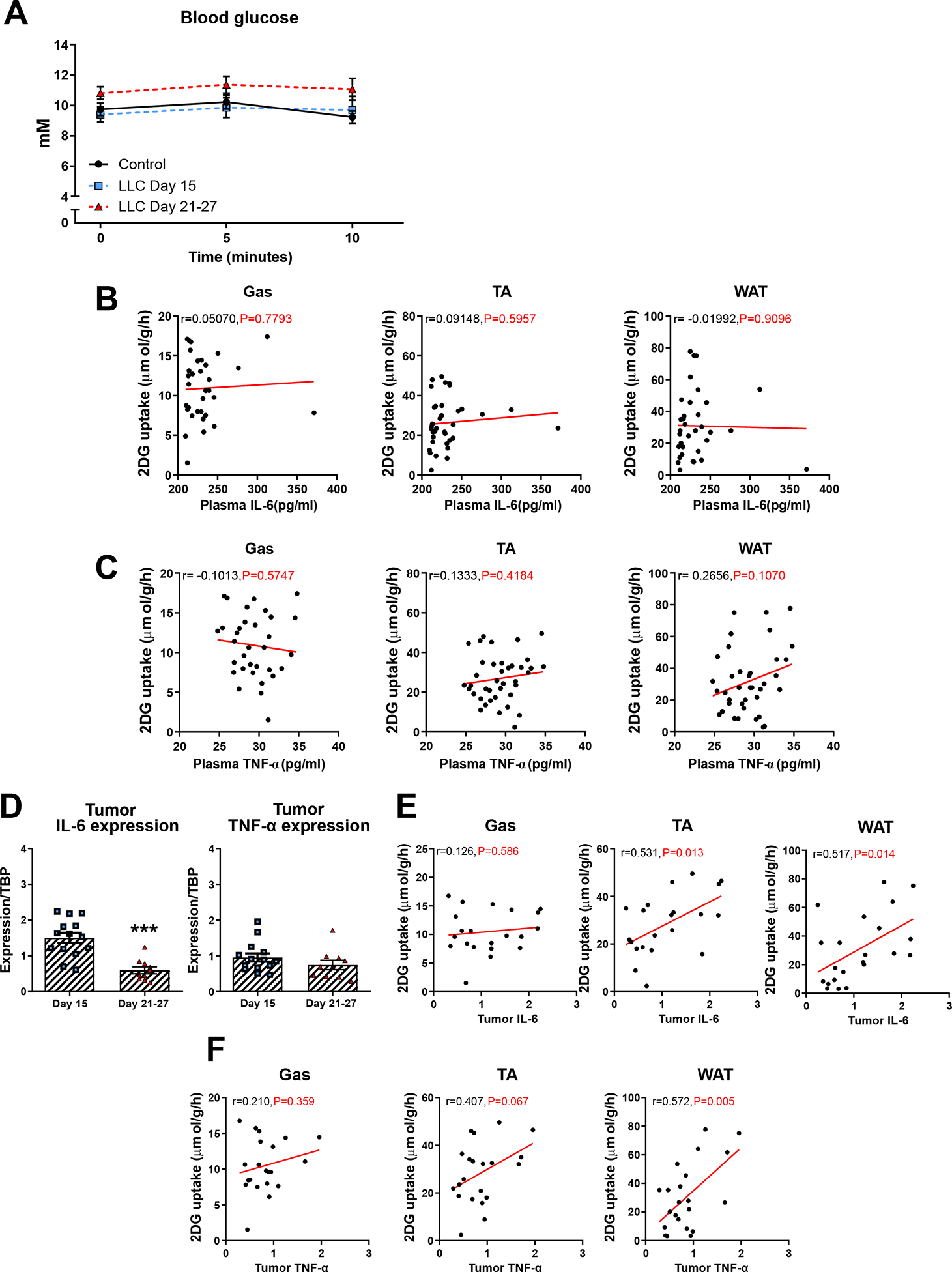
**A)** Blood glucose levels measured before (0 minutes), 5 minutes, and 10 minutes following retro-orbital saline injection in control (n=6) or Lewis lung carcinoma (LLC) tumor-bearing mice following 15 days (n=2) or 21-27 days (n=8) tumor inoculation. **B)** and **C)** Pearsons correlations between insulin-stimlated 2-DG uptake and plasma levels of interleukin-6 (IL-6) and tumor necrosis factor α (TNF-α) in gastrocnemius muscle (Gas), tibialis anterior muscle (TA), and white adipose tissue (WAT). **D)** mRNA expression of tumor tissue IL-6 and TNF-α. **E)** Pearsońs correlations between insulin-stimlated 2-DG uptake in peripheral tissues and tumor expression of IL-6. E) Pearsońs correlations between insulin-stimlated 2-DG uptake in peripheral tissues and tumor expression of TNF-α. Values are shown as mean± SE.

**Figure supplementary 2.**
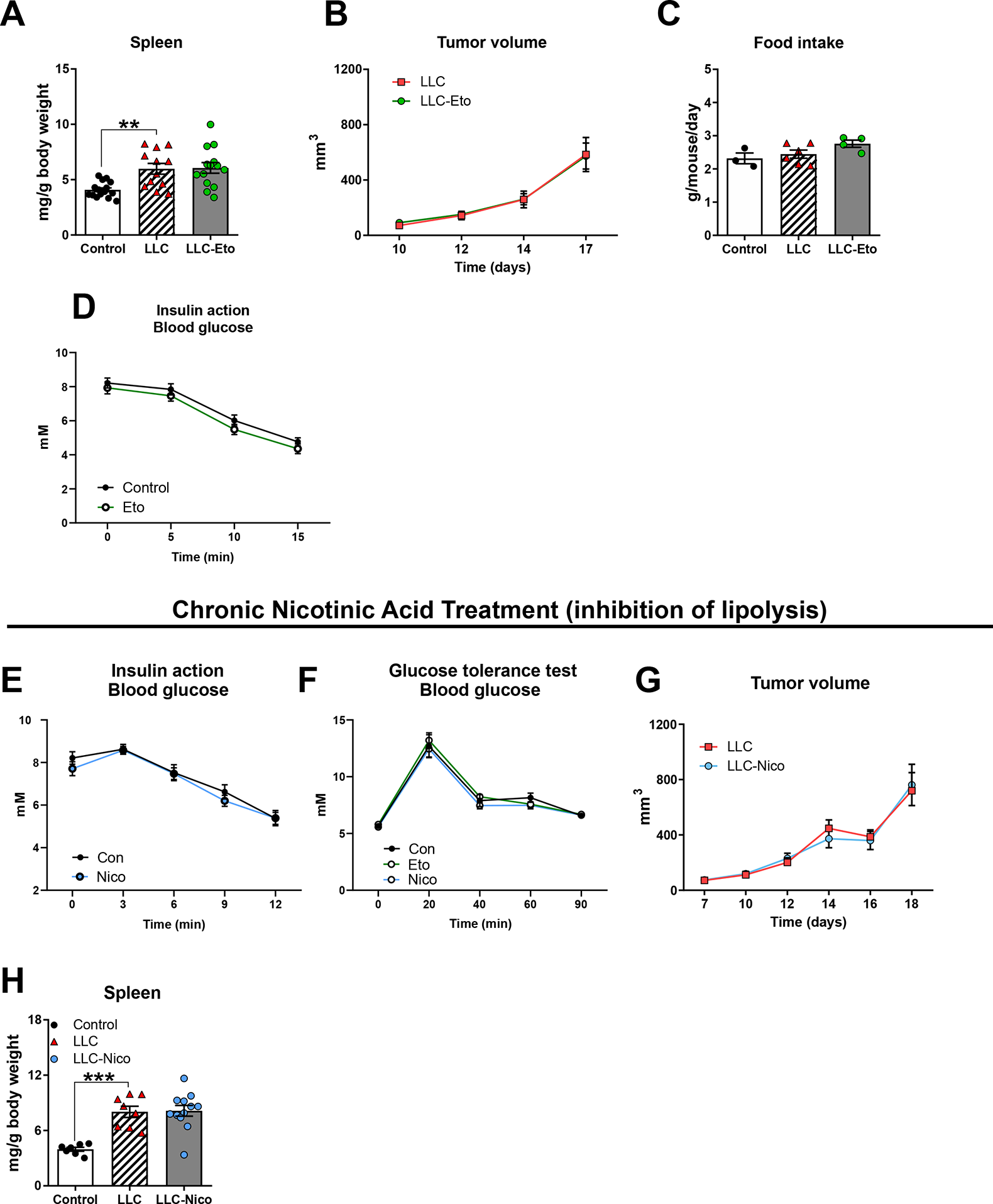
**A)** Spleen weight in control or LLC tumor-bearing mice following etomoxir (Eto) administration (n=11-14). **B)** Tumor volume of LLC tumor-bearing mice with Eto administration (n=11-14). **C)** Food intake in control or LLC tumor-bearing mice with Eto administration (n=3-6). **D)** Blood glucose concentration before (0 minutes), 5 minutes, 10 minutes and 15 minutes following retro-orbital insulin injection (0.3 U/kg body weight) in non-tumor-bearing control mice treated with or without Eto. **E)** Blood glucose concentration before (0 minutes), 5 minutes, 10 minutes and 15 minutes following retro-orbital insulin injection (0.3 U/kg body weight) in non-tumor-bearing control mice treated with or without Nicotinic acid (Nico). **F)** Blood glucose concentration before (0 minutes), 20 minutes, 40 minutes, 60 minutes and 90 minutes following intraperitoneal glucose tolerance test (GTT; 2 g kg-1 body weight) in control mice with or without Eto or Nico administration. **G)** Tumor volume and **H)** spleen weight in control or LLC tumor-bearing mice with Nico administration (n=8-12).Statistically significant effect of LLC is indicated by *P < 0.05; **P < 0.01; ***P < 0.001. Statistically significant effect of Nico in tumor-bearing mice is indicated by ##P < 0.01. Statistically significant effect of glucose injection on plasma insulin is indicated by $$$P < 0.001. Values are shown as mean ±SE with or without individual values.

**Supplementary Table 1.** Pearson’s correlations of tissue-specific glucose uptake, tumor volume, body weight, and time with (“Time”) or without (“none”) adjustment for time.

| Pearsons correlations |  |  |  |  |  |  |  |  |
| --- | --- | --- | --- | --- | --- | --- | --- | --- |
| Control | Variables |  | Glucose uptake_TA | Tumor volume | Glucose uptake_Gas | Glucose uptake_WAT | Body weight | Time |
| **-none<sup>a</sup>** | Glucose uptake_TA | Correlation | 1,000 | -,726 | ,463 | ,783 | ,013 | -,617 |
|  |  | p-value | . | ,000 | ,026 | ,000 | ,948 | ,001 |
|  |  | df | 0 | 24 | 21 | 21 | 24 | 24 |
|  | Tumor volume | Correlation | -,726 | 1,000 | -,401 | -,583 | ,042 | ,391 |
|  |  | p-value | ,000 | . | ,052 | ,003 | ,834 | ,044 |
|  |  | df | 24 | 0 | 22 | 22 | 25 | 25 |
|  | Glucose uptake_Gas | Correlation | ,463 | -,401 | 1,000 | ,281 | ,260 | -,033 |
|  |  | p-value | ,026 | ,052 | . | ,206 | ,221 | ,879 |
|  |  | df | 21 | 22 | 0 | 20 | 22 | 22 |
|  | Glucose uptake_WAT | Correlation | ,783 | -,583 | ,281 | 1,000 | ,013 | -,442 |
|  |  | p-value | ,000 | ,003 | ,206 | . | ,951 | ,031 |
|  |  | df | 21 | 22 | 20 | 0 | 22 | 22 |
|  | Body weight | Correlation | ,013 | ,042 | ,260 | ,013 | 1,000 | ,080 |
|  |  | p-value | ,948 | ,834 | ,221 | ,951 | . | ,693 |
|  |  | df | 24 | 25 | 22 | 22 | 0 | 25 |
|  | Time | Correlation | -,617 | ,391 | -,033 | -,442 | ,080 | 1,000 |
|  |  | p-value | ,001 | ,044 | ,879 | ,031 | ,693 | . |
|  |  | df | 24 | 25 | 22 | 22 | 25 | 0 |
| **Time** | Glucose uptake_TA | Correlation | 1,000 | -,669 | ,563 | ,723 | ,080 |  |
|  |  | p-value | . | ,000 | ,006 | ,000 | ,705 |  |
|  |  | df | 0 | 23 | 20 | 20 | 23 |  |
|  | Tumor volume | Correlation | -,669 | 1,000 | -,422 | -,497 | ,012 |  |
|  |  | p-value | ,000 | . | ,045 | ,016 | ,953 |  |
|  |  | df | 23 | 0 | 21 | 21 | 24 |  |
|  | Glucose uptake_Gas | Correlation | ,563 | -,422 | 1,000 | ,297 | ,263 |  |
|  |  | p-value | ,006 | ,045 | . | ,191 | ,225 |  |
|  |  | df | 20 | 21 | 0 | 19 | 21 |  |
|  | Glucose uptake_WAT | Correlation | ,723 | -,497 | ,297 | 1,000 | ,054 |  |
|  |  | p-value | ,000 | ,016 | ,191 | . | ,806 |  |
|  |  | df | 20 | 21 | 19 | 0 | 21 |  |
|  | Body weight | Correlation | ,080 | ,012 | ,263 | ,054 | 1,000 |  |
|  |  | p-value | ,705 | ,953 | ,225 | ,806 | . |  |
|  |  | df | 23 | 24 | 21 | 21 | 0 |  |

